## Supplementary Materials for "A causal test of affect processing bias in response to affect regulation"


### Supplemental Methods

Harmonized Preprocessing across Multiple Studies.

We preprocessed all imaging data using a single harmonized pipeline across raw MRI data collected for three separate studies – the current study, which we refer to as the Causal Test of Emotion Regulation (CTER) study (n=40), as well as two other previously published studies – the Intrinsic Neuromodulation of Core Affect (INCA) study[1] (n=19) and another study which augmented the INCA study with additional subjects (n=31; total n=50) that performed identical affect induction tasks to estimate the relative predictive fidelity of affective valence decoding models versus heart-rate deceleration, which we abbreviate as the HR study[2]. We reconstructed the decoding models for all subjects of all three studies using an identical image preprocessing and analysis pipeline. Preprocessing is described here.

All imaging data were acquired via a Philips 3T Achieva X-series MRI scanner (Philips Healthcare, Eindhoven, The Netherlands) with a 32-channel head coil. In brief, we acquired anatomic images with a MPRAGE sequence (matrix = 256 x 256, 220 sagittal slices, TR/TE/FA

= 8.0844/3.7010/8°, final resolution =0.94 x 0.94 x 1 mm$^3$. We acquired functional images with EPI sequence parameters: TR/TE/FA = 2000 ms/30 ms/90°, FOV = 240 x 240 mm, matrix = 80 x 80, 37 oblique slices, ascending sequential slice acquisition, slice thickness = 2.5 mm with 0.5 mm gap, final resolution 3.0 x 3.0 x 3.0 mm$^3$. All MRI data were processed using AFNI (Version AFNI_19.1.04)[3]. Anatomical data were processed via skull stripping, spatial normalization to the MNI152 brain atlas, and segmentation via FSL[4] into white matter (WM), gray matter (GM), and cerebrospinal fluid (CSF). Functional images underwent the following processing sequence: despiking; slice-time correction; deobliquing; motion correction; transformation to the spatially normalized anatomic image; regression of the mean time course and temporal derivative of the WM and CSF masks as well as a 24-parameter motion model[5,6], spatial smoothing (6-mm FWHM Gaussian kernel); and scaling to percent signal change. Gray matter (GM) masks for each subject were created using anatomical segmentation. A group-level GM mask was constructed, separately for each study, incorporating only GM voxels present in ≥ 50% of the individual subject GM masks within that study.

<u>Comparing Neural Encodings of Affective Image Stimuli across Multiple Studies</u>.

We computed study-specific group-level encodings from fMRI data according to the methods described in the main manuscript (see Materials and Methods: Affect Processing State Encodings), restricting those encodings to the GM masks computed for each separate study. We then conducted comparisons, separately for the CTER study against the HR study and the CTER study against the INCA study. The rationale for these comparisons was that the HR and INCA studies validated the predictive fidelity of the encoded brain states against independent psychophysiological measures of, respectively, valence (measured via heart rate deceleration) and arousal (measured via skin conductance response). Comparison of inter-study similarity of the encoding hyperplanes was conducted in two ways: (1) all gray matter voxels and (2) all gray matter voxels in which the encoding values survived global permutation testing ($p<0.05$). We then modeled the shared variance between the two encodings using linear regression in which the CTER study's encoding values were the independent variable and the comparison study's encoding values (either HR or INCA) were the dependent variable.

### Supplemental Results

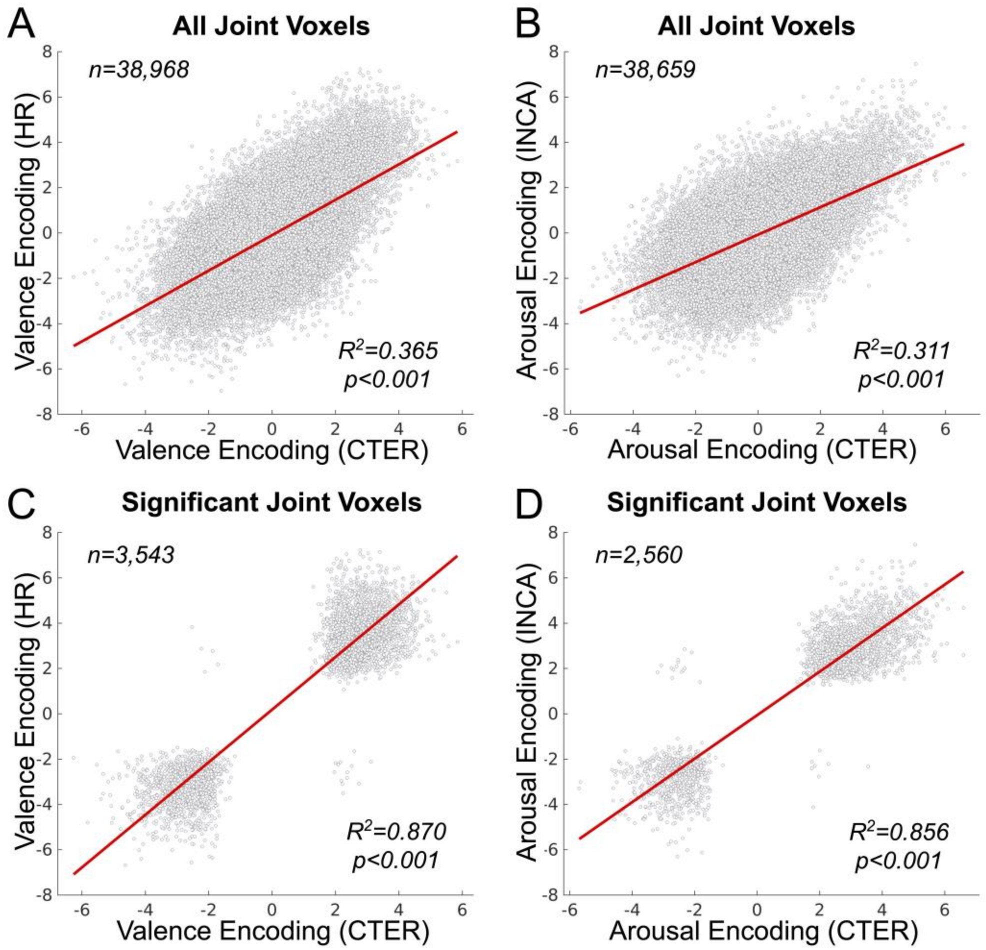


***Supplemental Figure S1:*** Comparing similarity of the derived neural encodings of affective image stimuli across multiple studies. ***(Top Row):*** *Inter-study comparison that includes the encoding values of all joint GM voxels shown for (A) affective valence processing and (B) affective arousal processing.* ***(Bottom Row):*** *Inter-study comparison that include encoding values for only*

*those joint GM voxels that survive global permutation significance testing (p<.05) for (C) affective valence processing and (D) affective arousal processing. Voxel-wise relationships are depicted as gray circles. The regression fit of the voxel-wise relationships is represented by the bold red line in each subplot. Total surviving joint voxels for each comparison are provided in the top left of each subplot. Inter-study shared variance is provided in the bottom right of each subplot. P-values refer to the significance of the regression fit's linear coefficient (F-test, α=0.05).*

***Supplemental Table 1:*** *International Affective Picture Set Image Identification Numbers*

| Trial Type | Identification Numbers |
|---|---|
| Mod-PS | 8510, 9421, 3350, 7502, 9908, 3266, 3061, 6821, 9910, 3140, 2799, 2717, 8420, 7230, 3168, 2800, 8503, 4520, 9253, 5460, 9250, 5830, 4608, 8380, 9901, 2208, 2160, 9400, 7260, 5825, 8300, 4660, 4640, 8210, 9500, 9040, 7405, 9412, 2075, 3131 |
| Mod-FS | 8034, 9419, 2071, 8490, 9570, 9300, 5480, 3301, 9830, 8170, 4680, 3215, 9183, 8370, 3225, 9921, 3064, 4599, 7350, 5450 |

**Supplemental Source Code and Data Availability.**

The authors have made the source code used in this supplemental analysis publicly available: https://github.com/kabush/CMPhaufe. The authors have also made the derivative activation maps necessary to execute this code available via the project's Open Science Framework Repository: https://osf.io/yn4vq/.

**Supplemental References**


1. Bush KA, Privratsky A, Gardner J, Zielinski MJ, Kilts CD. Common Functional Brain States Encode both Perceived Emotion and the Psychophysiological Response to Affective Stimuli.

Scientific Reports [Internet]. 2018 Dec [cited 2018 Oct 18];8(1). Available from: http://www.nature.com/articles/s41598-018-33621-6
